# Loss of replication and transcription systems accompanying transition to nucleus-dependent replication in “Ariadnavirales”, a proposed new order in nucleocytoviricot class *Megaviricetes*

**DOI:** 10.64898/2026.08.29.747986

**Authors:** Natalya Yutin, Yuri I. Wolf, Mart Krupovic, Eugene V. Koonin

## Abstract

*Sicyoidochytrium minutum* DNA virus (SmDNAV) was isolated several years ago from a protist host of family Thraustochytriaceae *of the* class Labyrinthulomycetes. This virus shared little similarity to other viruses in gene content and protein sequences, albeit seemingly belonging to the phylum *Nucleocytoviricota*. By extensive searches in genomic and metagenomic sequence databases, we identified numerous long contigs related to the SmDNAV genome and analyzed proteins shared by these putative viruses. Phylogenetic analyses place these viruses within the class *Megaviricetes*, outside of all established orders, and as a sister group to the clade combining families *Mamonoviridae* and ”*Manesviridae*”. Homologs of SmDNAV proteins were found in association (either integrated or co-sequenced) with other Labyrinthulomycetes and Rhodophyta protists from diverse marine and freshwater environments. Consequently, we propose SmDNAV as the prototype member of a new order, provisionally named “Ariadnavirales”, within class *Megaviricetes*, phylum *Nucleocytoviricota*. Members of “Ariadnavirales” have lost most of the genes encoding components of the replication and transcription systems that are otherwise conserved in nucleocytoviricots, suggestive of transition to genome replication and expression dependent on the host nucleus.

**Importance:** The viral phylum *Nucleocytoviricota* consists of diverse viruses with double-stranded DNA genomes within a broad size range from about 40 kilobases to about 5 megabases. The great majority of nucleocytoviricots encode DNA and RNA polymerases, and other proteins involved in genome replication and expression, and are known or predicted to replicate within viral ‘factories’ in the host cell cytoplasm. In recent years, several new groups of nucleocytoviricots have been discovered, primarily, by metagenome mining. Here we describe a putative new order of nucleocytoviricots, which we provisionally name “Ariadnavirales” for their protist hosts. These viruses have lost most of the genes responsible for genome replication and expression in other nucleocytoviricots, and their replication apparently depends on the host nucleus.

## Introduction

The phylum *Nucleocytoviricota* in kingdom *Bamfordvirae* of realm *Varidnaviria* unites diverse eukaryotic viruses with double-stranded (ds) DNA genomes that range in size from about 40 kilobases (kb) to about 5 megabases (mb) (1–6). The phylum currently consists of 3 classes: *Megaviricetes,* with orders *Imitervirales, Pimascovirales* and *Algavirales*, that encompass most of the nucleocytoviricot diversity, including viruses with the largest genomes; *Pokkesviricetes*, with orders *Asfuvirales* and *Chitovirales*, and putative order “*Egovirales”*, including a smaller set of viruses with moderate-size genomes, in particular, *Poxviridae*, by far the best studied group of nucleocytoviricots; and *Mriyaviricetes*, a recently identified group of viruses with the smallest genomes within the phylum (about 40 kb)(4, 7–9). The nucleocytoviricots share a common ancestry as evidenced by the conservation of about 20 core genes in the great majority of the phylum members (1, 3, 10). These core genes encode proteins involved in virion morphogenesis, including the major capsid protein (MCP) and genome packaging ATPase; key proteins mediating viral genomes replication, such as family B DNA polymerase (DNAP) and helicase-primase; and transcription, such as multiple RNA polymerase (RNAP) subunits and several transcription factors.

Most of the nucleocytoviricots that have been studied experimentally, in particular, vaccinia virus, a classic model of molecular virology, replicate in cytoplasmic viral factories, distinct non-membrane-bounded compartments (5, 6, 11–13). In these factories, viruses have no access to the host replication and transcription machineries, and rely virus-encoded protein toolkits for both processes. As suggested by both the comparative-genomic, namely, the conservation of the suites of replication and transcription proteins, and the experimental evidence (1), cytoplasmic replication is likely to be ancestral in the great majority of nucleocytoviricots, with the exception of the earliest-branching mriyaviricetes that lack most of these proteins (14). However, several other groups of nucleocytoviricots, such as those in families *Iridoviridae*, *Phycodnaviridae*, and “*Mininucleoviridae*”, have lost some components of the replication and transcription machineries, such as RNAP subunits, and have been shown or predicted to have fully or partially shifted to nuclear replication and expression (15, 16)(6, 17). Furthermore, a distinct nucleus-dependent replication strategy has been described for members of the proposed family ”*Manesviridae*”, involving the breakdown of the nuclear membrane followed by the packaging of nascent virions directly within the nucleoplasm (18).

Lately, the enormous, still incompletely sampled diversity of nucleocytoviricots has been expanding, primarily, through metagenome mining (19). Here, we employ this approach to describe a putative new nucleocytoviricot order that includes viruses with genomes of about 200 kb that display the most extensive, near complete loss of the replication and transcription machineries. The prototype of this putative viral order is the Sicyoidochytrium minutum DNA virus 001 (SmDNAV) (20, 21), for which ‘nuclear disappearance’ has been described, similar to manesviruses, whereas the rest of the members were discovered in metagenomes. We show that these viruses form a deep clade in the nucleocytoviricot class *Megaviricetes* and provisionally name the new order “Ariadnavirales”, after the mythical Cretan princess Ariadna who helped the hero Theseus to escape the deadly Knossos Labyrinth – recognizing the protist class Labyrinthulomycetes that includes *Sicyoidochytrium minutum*, the host of the prototype virus in the new order, but also acknowledging the likely broader host range of these viruses.

## Results

### Nucleocytoviricots in the putative order “Ariadnavirales” lost most of the genes involved in genome replication and transcription

Searching GenBank and IMG/VR protein and nucleotide sequence databases with the amino acid sequence of SmDNAV major capsid protein (MCP) as a query, we identified numerous related MCPs from various sources (Supplementary Table 1). In the phylogenetic tree of nucleocytoviricot MCPs, these proteins formed a distinct, strongly supported clade including a subclade consisting of SmDNAV and its close relatives (Figure 1 and Supplementary Figure S1). Among the contigs encoding SmDNAV-like MCP, there were two with the lengths close to that of the SmDNAV genome (over 200 kbp), potentially corresponding to complete or nearly complete virus genomes. These two contigs were sequenced as part of the cellular genomes of *Durusdinium trenchii* (SAR supergroup, Alveolata, Dinophyceae; accession #: CAXAMM010014914.1) and *Aurantiochytrium* sp. 8W (SAR supergoup, Stramenopiles; Bigyra; Labyrinthulomycetes; accession #: BAAFGW010000056.1). Searching the translations of the *D. trenchii* and *Aurantiochytrium* contigs with the SmDNAV proteins revealed 92 and 44 homologs, respectively. These contigs contained direct terminal repeats and hence were considered to represent complete circular genomes of SmDNAV-related viruses.

**Figure 1.**
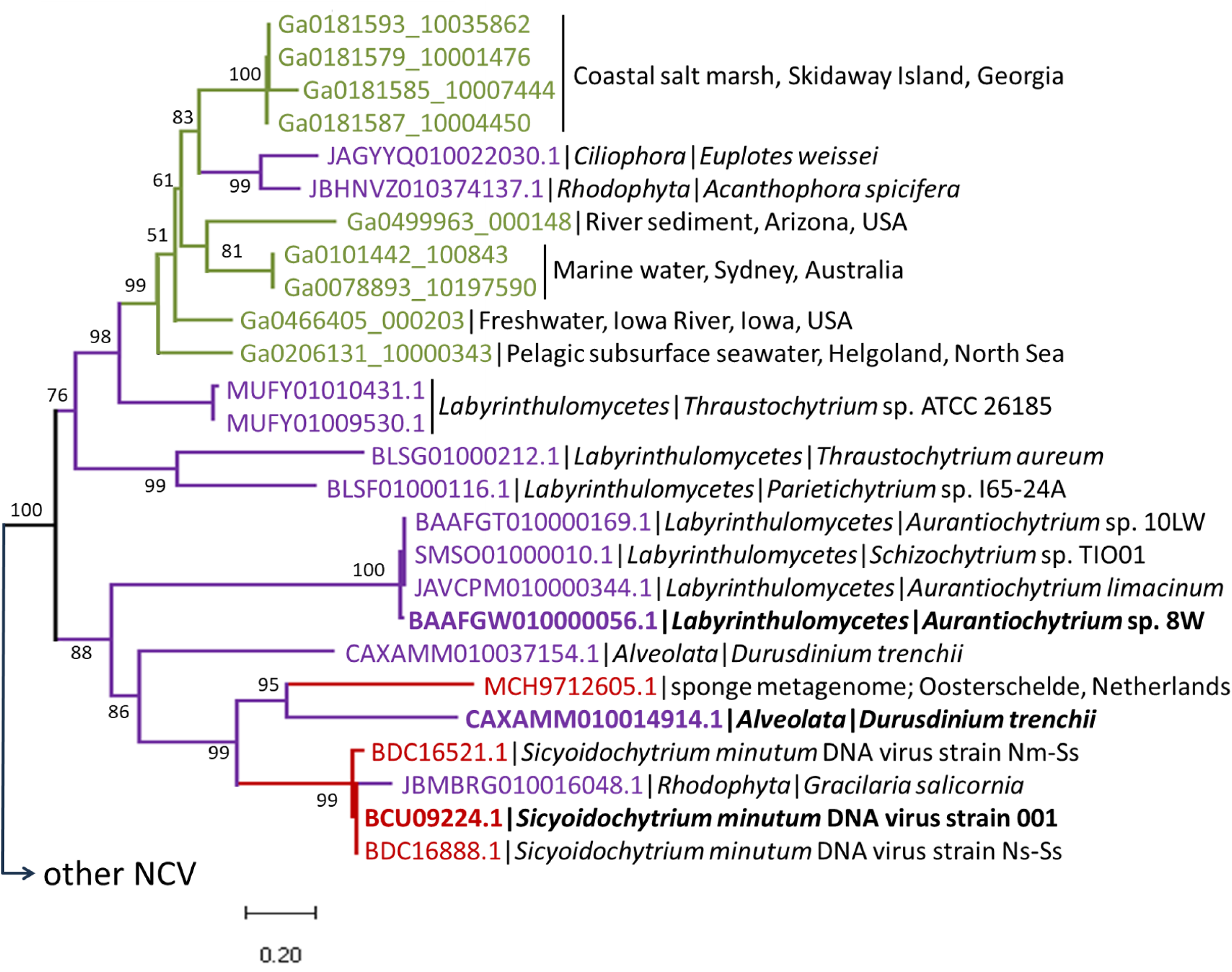
Phylogenetic tree of major capsid proteins of SmDNAV and related “ariadnaviruses”. Branch and sequence ID colors are according to the source: green, JGI IMG VR; purple, NCBI protist genome assemblies; red, NCBI GenBank. MUSCLE5 alignment; FastTree with WAG evolutionary model and gamma-distributed site rates.

Searching the GenBank protein sequence database with other SmDNAV proteins as queries, we identified multiple significant hits to protein sequences encoded in different bacterial metagenome assembled genomes (MAGs) (Supplementary Figure S2). The largest number of hits was to the MAG annotated as *Flavobacteriaceae* bacterium sequenced as part of the microbiome associated with the foraminifera *Cassidulina limbate* (SAR supergroup, Rhizaria, Foraminifera, Globothalamea) (22). Six contigs from this assembly (JAAEPP010000305.1, JAAEPP010000213.1, JAAEPP010000310.1, JAAEPP010000199.1, JAAEPP010000385.1, and JAAEPP010000770.1; total length: 139,814 bp; contig length range: 11,890 to 33,908 bp; 52 homologs of SmDNAV proteins). In all likelihood, the six contigs represent a partial genome of an SmDNAV-like virus associated with *C. limbate and* misannotated as bacterial. Thus, the six contigs and the two apparently complete genomes of SmDNAV-like viruses were subject to further analysis. The proteins encoded in these contigs and SmDNAV were clustered by sequence similarity, yielding 60 clusters containing three or more proteins each (Supplementary Table 1; https://ftp.ncbi.nih.gov/pub/yutinn/lbr_2026/). Sequences within these clusters were aligned and used as queries in a translating search for additional homologs of the SmDNAV proteins in the JGI IMG/VR database and NCBI protist genome assemblies. The retrieved contigs were translated, and the search with SmDNAV queries was repeated against this set to link the new proteins to their SmDNAV homologs. Finally, approximate phylogenetic trees were constructed to identify proteins forming clades with SmDNAV homologs. (Supplementary Table 1 and https://ftp.ncbi.nih.gov/pub/yutinn/lbr_2026/). SmDNAV and SmDNAV-like viruses associated with *D. trenchii* and *Aurantiochytrium* sp. 8W were predicted to encode 358, 240 and 231 proteins, respectively, yielding estimated gene densities of 1.5, 1.1 and 0.9 gene/kilobase, values typical of nucleocytoviricots (1) except for the first one that appeared to include over-prediction of short proteins.

The 60 clusters of homologous proteins from the group of viruses related to SmDNAV (hereafter ariadnaviruses) were annotated by searching Conserved Domain Database (CDD), Pfam and PDB databases (see Methods), and also by comparing the predicted protein sequences to NCVOGs (nucleocytoviricots orthologous groups) (23), to link them to orthologs in other nucleocytoviricots.

Ariadnaviruses encode the nucleocytoviricot core structural and morphogenetic proteins, namely, MCP, packaging ATPase and disulfide oxidoreductase of the Erv1/Alr family as well as the papain-like thiol protease of the OTU family shared with nucleocytoviricots of order *Pimascovirales* and some groups of *Algavirales* (Supplementary Table 2, Figure 2). The OTU protease is only distantly related to the papain-like protease involved in the capsid protein maturation in other nucleocytoviricots (prototyped by vaccinia virus I7) (24), but given the absence of the latter in the OTU-encoding viruses, can be tentatively implicated in the same function. In addition, ariadnaviruses encode a homolog of the chlorovirus PBCV-1 P4, a minor structural protein shown to stabilize the interactions between neighboring MCP capsomers (25). Although no homologs of the entry-fusion complex (EFC) conserved in most nucleocytoviricots were initially annotated in SmDNAV, we identified three paralogs of disulfide-bonded proteins (vaccinia virus A16/G9/J5 homologs) that in poxviruses are subunits of the EFC (26, 27) and are scattered among nucleocytoviricots (Figure 2), and one protein homologous to the disulfide-bonded, myristoylated virion proteins L1/F9. The apparent closest relatives of these ariadnavirus proteins were identified in iridoviruses (Supplementary Figures S3, S4). The conservation of L1R/F9L in most nucleocytoviricots correlates with the conservation of the thiol-disulfide oxidoreductase, the ortholog of vaccinia virus E10R, which is required for the formation of disulfide bonds in L1R and F9L (28).

**Figure 2.**
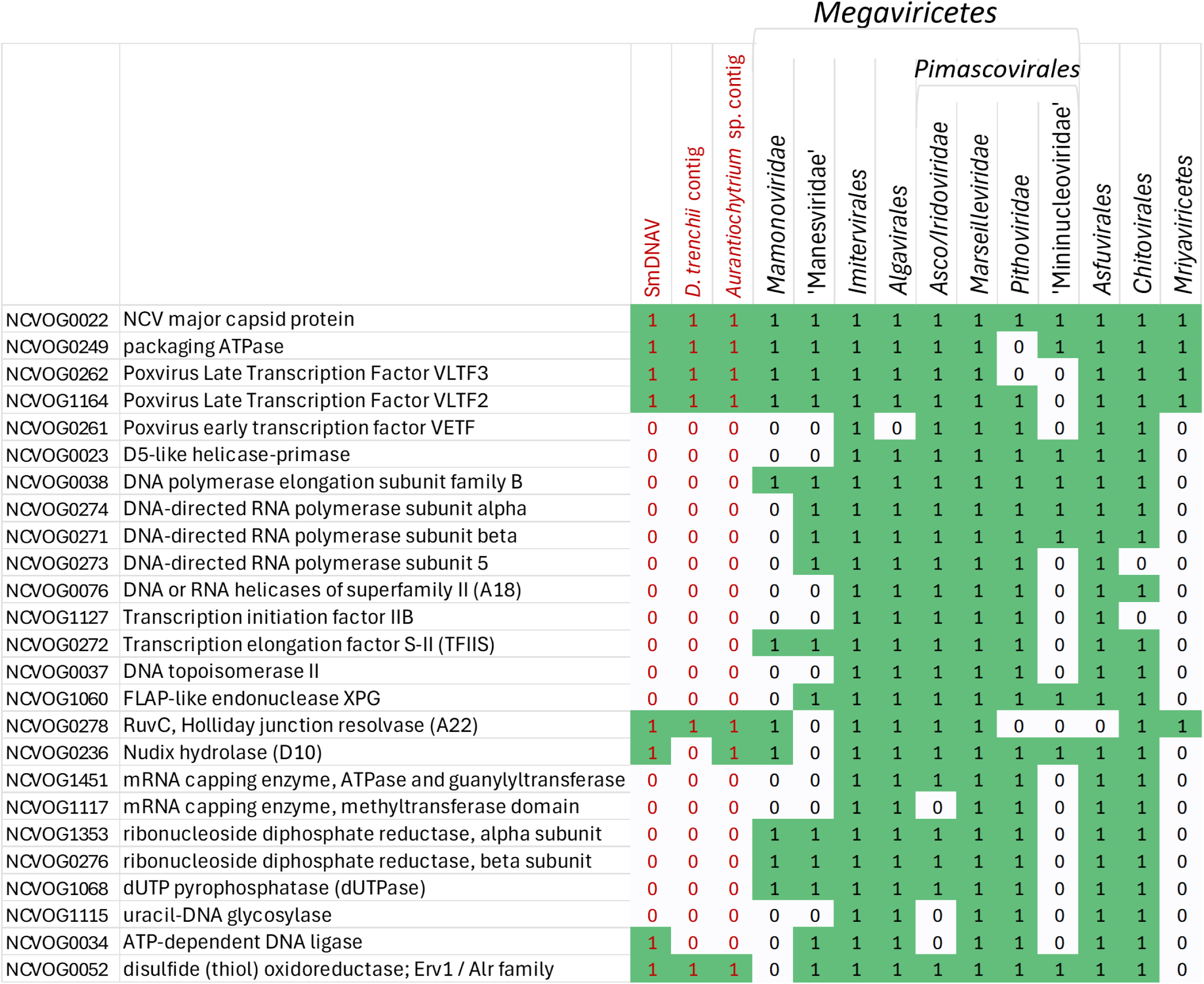
Presence-absence of core nucleocytoviricot proteins across major groups within the phylum. ‘1’ represents presence and ‘0’ represents absence of the respective gene.

By contrast, the majority of the core genes encoding proteins involved in genome replication and transcription of nucleocytoviricots, including DNAP, D5-like primase-helicase, other helicases, RNAP, some eukaryotic transcription factors and the two subunits of the mRNA capping enzyme, are missing in ariadnaviruses (Figure 2 and Supplementary Table 2). The only proteins involved in information processing present in ariadnaviruses are a homologs of the eukaryotic transcription factor TFIIB, two nucleocytoviricot-specific transcription factors (VLTF2 and VLTF3), YqaJ-like viral recombinase, Holiday junction resolvase (RuvC homolog), and Proliferating Cell Nuclear Antigen (PCNA), the sliding clamp replisome subunit (Figure 2 and Supplementary Table 2). In addition, SmDNAV but not two other complete ariadnavirus genomes encode an ATP-dependent DNA ligase shared with many nucleocytoviricots (Figure 2). SmDNAV also encodes an AAA+ ATPase related to the origin recognition complex subunits conserved in archaea and eukaryotes (SmDNAV_BCU09089). Finally, SmDNAV encodes a Pif1-like superfamily I helicase and a distinct primase-helicase related to the origin-binding UL9-like proteins conserved in orthoherpesviruses and malacoherpesviruses (29). The latter SmDNAV protein contains an N-terminal archaeal-eukaryotic primase (AEP) superfamily primase-polymerase domain (270-515 aa) and a C-terminal superfamily II helicase domain (788–1409).

Related proteins are also encoded by mimiviruses (R8 protein (30)), African swine fever virus (pF1055L) and a subset of mriyaviruses (14). Notably, whereas in orthoherpesviruses, the primase domain is inactivated, the catalytic AEP motifs are intact in the SmDNAV protein, similar to homologs from nucleocytoviricots and malacoherpesviruses. Apart from these highly conserved proteins, we also identified a diverged homolog of the OB-fold single-stranded DNA-binding proteins that is shared with many groups of nucleocytoviricots (Supplementary Figure S6, Supplementary Table 2) and T7-like bacteriophages (31). Among nucleocytoviricots, only viruses of the class *Mriyaviricetes* that have much smaller genomes are depleted of genes involved in replication and transcription to a similar extent as ariadnaviruses (Figure 2) (14). Some of these genes are missing also in families *Mamonoviridae* and “*Manesviridae*” but viruses of both families encode a DNAP and members of “*Manesviridae*” encode RNAP subunits as well (18) (Figure 2).

To investigate the relationships between ariadnaviruses and other nucleocytoviricots, we constructed a phylogenetic tree from a concatenated alignment of three core proteins that are conserved in nearly all nucleocytoviricots, namely, MCP, packaging ATPase and transcription factor VLTF3. In this tree, ariadnaviruses are confidently placed within *Megaviricetes* as the sister group to the clade consisting of *Mamonoviridae* and “*Manesviridae*” (Figure 3). Considering the relative evolutionary depth of the branch joining *Mamonoviridae*-“*Manesviridae*” with ariadnaviruses (Supplementary Figure S6), we propose that the latter should be classified as a new order “Ariadnavirales”, and the former two families become another order as recently proposed (18), which can be tentatively named “*Manovirales*” (after MAnesviridae and MamoNOviridae). Given that the clade consisting of the two putative new orders is confidently included within the expansive megaviricetes clade (Figure 3), all the rest of which encode full complements of replication and transcription system components, it is effectively certain that these genes were lost. Most of these losses apparently occurred at the branch leading to the common ancestor of the putative new orders, and some additional genes were lost in “Ariadnavirales”.

**Figure 3.**
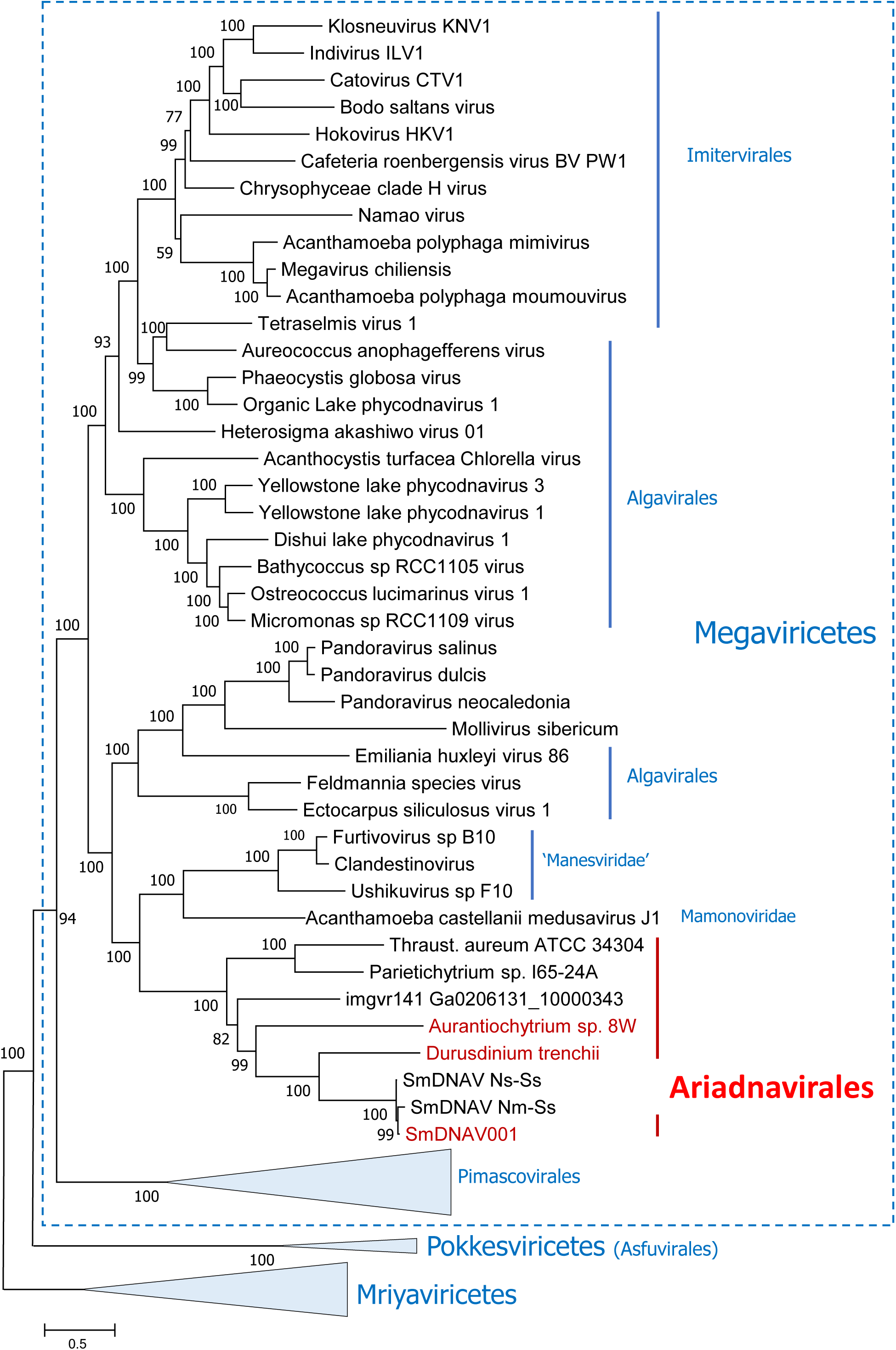
Phylogenetic tree of nucleocytoviricots. The tree was constructed from concatenated MUSCLE5 alignments of MCP, packaging ATPase and VLTF3. IQ-Tree, Q.pfam+F+R7 evolutionary model; aBayes support values are indicated.

### Non-core genes of “Ariadnavirales”

Apart from the conserved core proteins, nucleocytoviricots with larger genomes encode numerous proteins that are known to be involved or are implicated in lineage-specific morphogenetic, replication and expression processes, and various virus-host interactions (6). Our annotation pipeline (see Methods) yielded prediction of conserved domains and biochemical functions for 53 proteins that were denoted hypothetical in the original annotation of the SmDNAV genome (21) (Supplementary Table 2).

Given that ariadnaviruses are the sister group of manoviruses in the phylogenetic tree of hallmark proteins (Figure 3), and further, that at least one ariadnavirus shares the host with a mirusvirus (32, 33), we searched for proteins that might be specifically shared between ariadnaviruses and these two groups of viruses. Unexpectedly, given the strongly supported phylogenetic grouping, we failed to identify any derived shared characters (synapomorphies) of the ariadna-manovirus clade – that is, proteins shared by these viruses to the exclusion of other nucleocytoviricots. Nevertheless, examination of several shared genes appears instructive (Supplementary table S2). A prominent trend is the apparent convergent acquisition of many if not most of the shared genes, a phenomenon that is increasingly recognized as central to viral evolution (4, 34). Notable examples include high mobility group (HMG)-box domain protein implicated in chromatin structuring that is present in ariadnaviruses and medusaviruses but apparently were captured from eukaryotic hosts independently; HNH family endonuclease that is conserved in ariadnaviruses and in manesviruses but appear to have been independently acquired from different phages; and Tlr 6Fp, an uncharacterized protein encoded by ariadnaviruses and manesviruses, but in the former case, grouping with homologs from transpovirons and virophages, and in the latter case, with prasinoviruses; and some additional, uncharacterized proteins.

Several non-core genes labyrinthoviruses shared with mirusviruses were identified as well (Supplementary table S2). Homologs of a divergent serine/threonine kinase of SmDNAV (BCU09292) were detected in mirusviruses (YAS67345, YAS67549, YAS67601), but phylogenetic analysis showed that these kinases likely originated from different branches of eukaryotes.

BCU09408 protein of SmDNAV and Mirusvirus proteins YAS67408 are RecD-like helicases that belong to the same clade, together with some eukaryotic homologs including those from labyrinthulomycetes *Hondaea fermentalgiana* and *D. trenchii*. This case might reflect a network of horizontal gene transfers among ariadnaviruses, mirusviruses and their protist hosts.

SmDNAV protein BCU09335 and Mirusvirus YAS67499 are S8 family peptidases that, once again, appear to have different origins in ariadnaviruses and mirusviruses.

Both ariadnaviruses and mirusviruses encode heliorhodopsins (SmDNAV protein BCU09291) that also show a scattered presence among nucleocytoviricots (35, 36) and appear to have multiple, independent origins. Heliorhodopsins are light-dependent proton transporters that can depolarize the host membrane, potentially, inactivating host defenses and/or preventing superinfection (36).

A more complex case is a protein family represented by SmDNAV Nm-Ss protein BDC16459. Full-length homologs were identified in *D. trenchii* and *Ariadnamicetes* contigs, as well as in mirusvirus proteins. This protein consists of two domains, of which the N-terminal domain is much stronger conserved than the C-terminal one. The N-terminal domain of BDC16459 and its homologs is distantly but significantly similar to 7P0U_F, an apoptosis inhibitor of Orf virus (Poxviridae). The C-terminal domain shows distant but significant similarity to NucS endonuclease (pfam01939) that cleaves ssDNA regions of branched DNA structures and is represented by multiple paralogs across ariadnaviruses. In SmDNAV001, BCU09076 shares the NucS endonuclease C-terminal domain with SmDNAV protein BDC16459, but contains an unrelated N-terminal domain; another variant of this domain is encoded by a distinct SmDNAV001 gene. This domain is widespread in phaeoviruses and their hosts. It shows strong similarity to the N-terminal domain of the P22 phage antirepressor protein (pfam10547) and to the putative MSV199 domain-containing protein of Invertebrate iridescent virus 6.

Ariadnaviruses also share with mirusviruses an uncharacterized protein (SmDNAV BCU09119 and mirusvirus YAS67358) that belongs to an expansive family represented in some nucleocytoviricots, bacteria and eukaryotes. Phylogenetic analysis places these ariadnavirus and mirusvirus proteins in a clade with homologs from “Manesviridae”, and some bacteria, suggestive of multiple HGT events.

A notable feature of the ariadnavirus proteome is the enrichment in carbohydrate metabolism proteins (Supplementary Table S2). For instance, SmDNAV encodes 15 proteins from this functional category, including multiple glycosyltransferases and glycoside hydrolases of different families as well as sugar transporter of the SWEET family. Multiple glycosyltransferases are encoded also by some other nucleocytoviricots, in particular, chloroviruses and mimiviruses, and are involved in glycosylation of virion proteins (37). The abundance of glycosyltransferases in ariadnaviruses implies that some of their virion proteins encompass complex glycans. By contrast, glycoside hydrolases (n=3 in SmDNAV) could be involved either in carbohydrate biosynthesis or aid during virion entry and/or egress. For instance, SmDNAV ORF003 (BCU09069) encodes a putative chitinase, which could locally degrade the cell wall of the host, enabling the virus to reach the cell membrane, as has been proposed for chloroviruses (38). In addition, SmDNAV encodes 5 proteases, 2 DNA Methyltransferases, 2 poly (ADP-ribose) polymerases, a diadenosine tetraphosphate (Ap4A) hydrolase and several distinct nucleases, which are all likely involved in virus-host interaction, in particular, counterdefense (Supplementary table S2).

## Discussion

The discovery of new groups of viruses by genome and, above all, metagenome analysis not only steadily expands the virosphere but delivers indications of new biology. The putative order “Ariadnavirales” described here is notable for the unprecedented, near complete loss of genes encoding components of the replication and transcription systems while retaining a relatively large genome size of about 200 kb that is typical of nucleocytoviricots. Most of these proteins are missing also in “Manovirales”, the sister group of “Ariadnavirales”, suggesting an ancient gene loss that, in all likelihood, accompanied transition to nuclear replication. Indeed, replication within the nucleus has been demonstrated for several members of “Manovirales” (18, 39–41).

However, other manovirals, such as ushikuvirus, while apparently replicating in the host cytoplasm, have been reported to disrupt the nuclear membrane, likely, recruiting nuclear proteins to their cytoplasmic factories (41). Similarly, cytoplasmic replication and disappearance of the nuclear membrane have been observed in SmDNAV-infected cells (20). Thus, ariadnavirals apparently have not switched to nuclear replication but rather evolved a distinct mode of hijacking the nucleus while retaining cytoplasmic replication. Notably, manoviruses encode the full set of 5 histones that form nucleosome structures on the viral genomic DNA (42–44). Ariadnaviruses lack histones but instead encode an HMG protein, suggesting an alternative mode of viral chromatin compactization.

Apart from “Manovirales” and “Ariadnavirales”, two groups of nucleocytoviricots lack genes for most components of the replication and transcription machineries, the proposed family “Mininucleoviridae” in the order *Pimascovirales* (17, 45) and the class *Mriyaviricetes* (14). The viruses in these groups have the smallest genomes among the nuclocytoviricots, 70-80 kb and 35-40 kb, respectively. mininucleovirids have been shown to replicate in the host cell nucleus (17). Although the information on the replication of yaravirus, the only cultivated member of the *Mriyaviricetes*, remain scarce, this virus appears to form viral factories in the cytoplasm, in the area previously occupied by the nucleus (8), suggesting a similar strategy of nuclear dissolution induced ariadnavirals and manesvirids (18, 20, 41). Despite adopting similar replication strategies, the three groups of viruses appear to have evolved along completely different evolutionary trajectories. Mininucleovirids seem to represent the ultimate case of reductive evolution within the phylum, manifested not only by the extensive gene loss but also by accumulation of low-complexity sequences . By contrast, mriyaviricetes, the deepest branch of nucleocytoviricots, most likely, never possessed the replication and transcription machineries typical of this phylum (14). “Manovirales” and “Ariadnavirales” are notable in that their gene loss is highly selective, without general signatures of genome degradation.

A striking parallel with respect to the loss of the replication and transcription systems is apparent between “Manoviruses” and “Ariadnavirales” and several groups in phylum *Mirusviricota* that belongs to the second vast realm of viruses with large dsDNA genomes, *Duplodnaviria* (33, 46–48). Mirusviricots of the orders *Okeanovirales* and *Styxvirales* lack (nearly) all proteins involved in replication and transcription, and in parallel, have accumulated multiple spliceosomal introns, strongly suggestive of nuclear replication (48). No signs of intron invasion were detected in “Manovirales” and “Ariadnavirales”; the underlying causes of this striking difference between the evolutionary histories of the (predicted) nuclear nucleocytoviricots and mirusviricots remain enigmatic. One intriguing possibility is that, whereas at least some of the intron-rich mirusviricots cause persistent infection (32, 33), likely, preserving nuclear functions, conducive to both intron invasion and splicing, manoviruses and ariadnaviruses disrupt nuclear structure and functions, precluding intron acquisition.

Although “Manovirales” and “Ariadnavirales” comprise a strongly supported clade in the phylogeny of hallmark proteins, there are relatively few homologous genes in the two groups, apart from those of the nucleocytoviricot core. Most of the shared non-core genes appear to have been captured convergently, independently, by manovirals and ariadnavirals, suggestive of highly dynamic evolution in this clade of nucleocytoviricots.

A deeper foray into the evolution of the nucleocytoviricots reveals complex routes. Eukaryotic varidnavirians appear to have originally evolved from tectivirids (tailless phages) of phylum *Preplasmaviricota* whose immediate descendants in eukaryotes are polinton-like viruses replicating in the nucleus (49, 50). Given the basal position of *Mriyaviricetes* in the nucleocytoviricot phylogeny (14), it appears most likely that the last common ancestor of nucleocytoviricots has already transitioned to cytoplasmic replication (Figure 4). However, whereas mriyaviricetes remained dependent on the nuclear replication machinery, the last common ancestor of *Megaviricetes* and *Pokkesviricetes* apparently acquired the replication and transcription machineries from the host, gaining full autonomy and independence from the nucleus (Figure 4). During the subsequent 2 billion years or so of nucleocytoviricot evolution, several groups either returned to the nucleus or evolved alternative ways of exploiting the nucleus (Figure 4).

**Figure 4.**
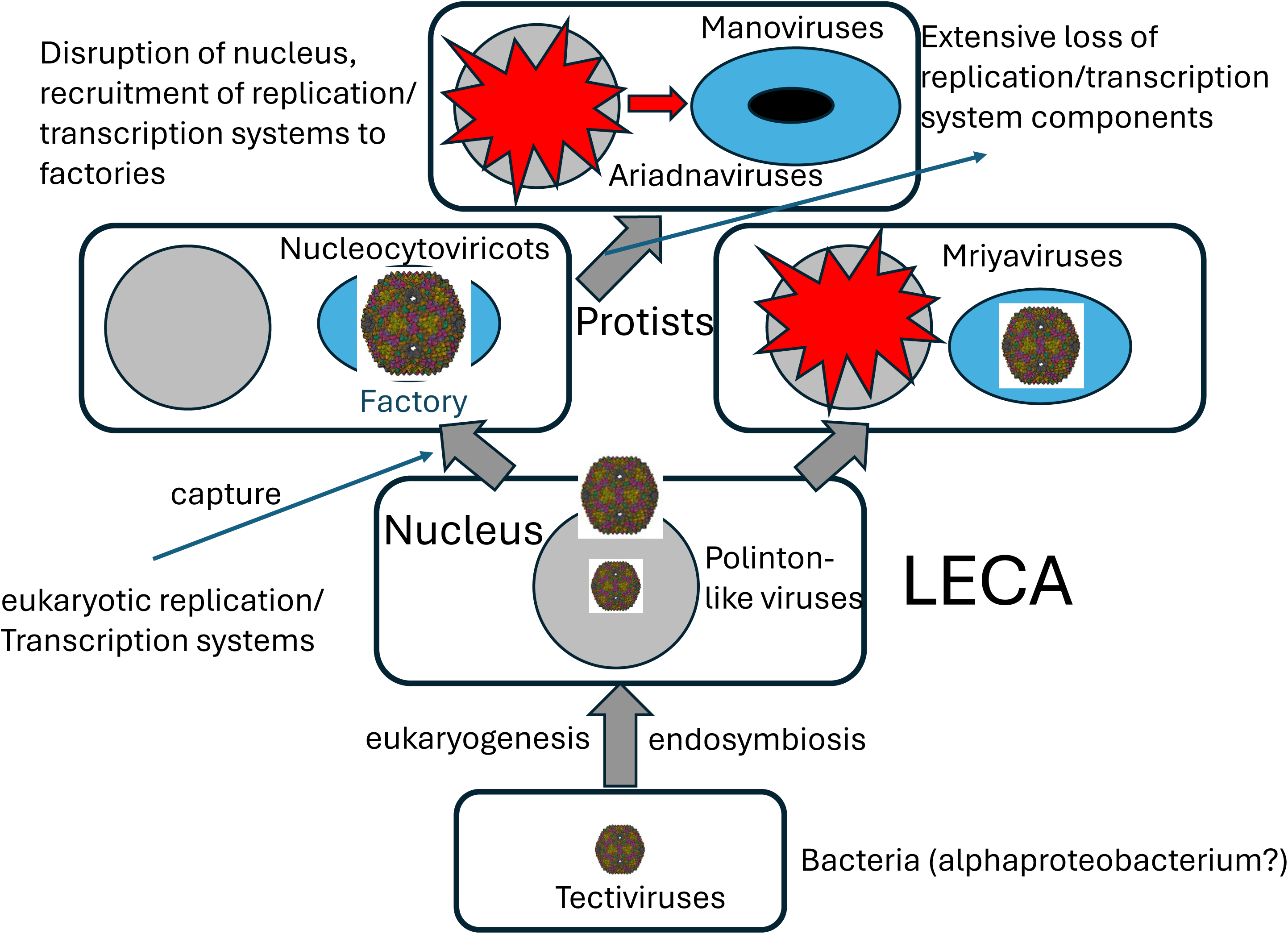
**Gain and loss of replication and transcription systems and inferred shuttling between nuclear and cytoplasmic replication sites in the evolution of nucleocytoviricots.** LECA, Last Eukaryotic Common Ancestor

While this manuscript was in preparation, a preprint describing a family of viruses related to SmDNAV (provisionally named “Skiaviridae”) and infecting thraustochytrid protists has been released (51).

## Conclusions

Combining genome and metagenome mining, we identified a putative new order of nucleocytoviricots, with SmDNAV as the only cultivated member and the prototype. These viruses have moderate-sized genomes of about 200 kb and are associated with a broad range of protists, including Stramenopiles, Alveolata, Rhizaria Ariadna and Rhodophyta, coming from various marine and freshwater environments. Phylogenetic analysis of core proteins shows that the proposed order “Ariadnavirales” belongs to class *Megaviricetes* as a sister group to another putative order, “Manovirales”, combining families *Mamonoviridae* and “Manesviridae”. The nucleocytoviricots in the “Ariadnavirales”-“Manovirales” have lost the genes encoding most components of the replication and transcription systems, indicative of dependence on the protein machinery for replication provided by the host nucleus as demonstrated for some member of “Manovirales”.

## Materials and Methods

### Clustering and annotation of ariadnavirus proteins

Initially, GenBank protein and nucleotide databases were searched with SmDNAV MCP as a query; candidate contigs of approximately 200 kbp in length were examined for the presence of other SmDNAV protein homologs. Two contigs, *Durusdinium trenchii* contig CAXAMM010014914.1 and *Aurantiochytrium* sp. 8W contig BAAFGW010000056.1, were found to have 92 and 44 SmDNAV-related proteins, respectively.

GenBank protein database searches with other SmDNAV proteins using BLASTP (52) revealed six MAGs ascribed to *Flavobacteriaceae* bacterium (JAAEPP010000305.1, JAAEPP010000213.1, JAAEPP010000310.1, JAAEPP010000199.1, JAAEPP010000385.1, and JAAEPP010000770.1), with significant presence of SmDNAV-related proteins (52 in total).

ORFs were predicted in *D. trenchii* contig CAXAMM010014914.1 using Prodigal (53), and combined with SmDNAV proteins, proteins encoded in *Aurantiochytrium* sp. 8W contig BAAFGW010000056.1, and the *Flavobacteriaceae* bacterium contigs. These proteins were clustered by finding reciprocal best hits with e-value below 10^-6^ across the contigs with subsequent manual refinement. Protein cluster sequences were aligned using MUSCLE 5 (54). The cluster alignments were compared to publicly available profile databases (PDB_mmCIF70_20_Feb, Pfam-A_v38.2, Uniprot-SwissProt-viral70_3, and NCBI_Conserved_Domains (CD)_v3.19) using HHPRED (55) (for cluster annotations, see Supplementary Table 2).

### Collecting ariadnavirus sequences from IMG/VR database and NCBI protist genome assemblies

For each ariadnavirus protein cluster alignment, a consensus sequence was calculated; the consensus sequences were used as queries in a translating search in the JGI IMG/VR database (56), and in NCBI protist genome assemblies (https://www.ncbi.nlm.nih.gov/genome/), retrieved using the Datasets tool (https://www.ncbi.nlm.nih.gov/datasets/docs/v2/command-line-tools/).

In the resulting contigs, ORFs were predicted using Prodigal in the metagenomic mode; SmDNAV proteins were searched against these ORFs; homologous proteins were collected and aligned using MUSCLE 5 with homologous proteins retrieved from Clustered nr database with SmDNAV proteins as queries. Approximate phylogenetic trees were constructed from the alignments, using Fasttree with WAG evolutionary model and gamma-distributed site rates (57). Ariadnavirus clades were identified in the trees as strongly supported branches containing a SmDNAV protein.

### Phylogenetic analysis

Multiple alignments of MCP, packaging ATPase, and VLTF3 were concatenated, the phylogenetic tree was built using IQ-TREE (58), with the Q.pfam+F+R7 evolutionary model, chosen according to BIC by the built-in model finder.

To establish the provenance of non-core ariadnavirus genes the following procedure was performed. First, ariadnavirus queries were used in a PSI-BLAST search (e-value threshold of 0.01) against the three databases: a collection of 47.5k completely sequenced prokaryotic genomes from NCBI Genome assemblies, EukProt database of curated eukaryotic proteins (59) and NCBI clustered NR database. Up to 1000 best hits were collected from each. Retrieved homologs were aligned with the ariadnavirus sequences and aligned using MUSCLE5. Approximate phylogenetic trees were reconstructed using FastTree with WAG evolutionary model and gamma-distributed site rates.

### Taxonomic depth of ariadnaviruses

The phylogenetic tree, constructed from the concatenated alignments of MCP, packaging ATPase and VLTF3, was ultrameterized by iteratively balancing the depths of the two subtrees at each internal node, preserving the total branch length. For each clade that (approximately) corresponded to a taxonomic rank (class, order, or family), the depths (distance from the leaves) were recorded for two key points: the last common ancestor of the taxon (the minimum taxon depth) and the bifurcation from sister taxa (the maximum taxon depth). Supplementary Figure S6 plots the depths of the taxa included in this tree, in comparison to ariadnaviruses.

### Protein structure prediction and analysis

3D structure of ariadnavirus protein clusters was predicted using ESM fold (60) on NIH HPC Biowulf cluster (https://hpc.nih.gov). Within the cluster alignments, the sequence most similar to the alignment consensus, was selected as the master sequence. Predicted structures were compared to the PDB structure database using FoldSeek (61).

## Data availability

This paper is based entirely on the analysis of existing, publicly available data. Data generated during downstream analysis are available in the Supplementary Material or via ftp at zenodo.doi or https://ftp.ncbi.nih.gov/pub/yutinn/lbr_2026. Any additional information required to reanalyze the data reported in this paper is available from the authors.

## Author contributions

N.Y. initiated the study, collected the data and performed research; N.Y., Y.I.W., M.K. and E.V.K. analyzed the data; E.V.K. wrote the manuscript that was read, edited and approved by all authors.

## Acknowledgements

The authors thank John Archibald for sharing data prior to publication and useful discussions. N. Y., Y.I.W., and E.V.K. are supported by the Intramural Research Program of the National Institutes of Health (NIH). This work utilized the computational resources of the NIH high-performance computing (HPC) Biowulf cluster (https://hpc.nih.gov). The contributions of the NIH author(s) are considered Works of the United States Government. The findings and conclusions presented in this paper are those of the author(s) and do not necessarily reflect the views of the NIH or the U.S. Department of Health and Human Services.

## Supplementary material

**Supplementary table S1**

Database search results for predicted ariadnavirus proteins

**Supplementary table S2**

Clusters of homologous ariadnavirus proteins

**Supplementary figure S1**

Phylogenetic tree of the major capsid proteins of “Ariadnavirales”

**Supplementary figure S2**

Provenance of ariadnavirus MCP sequeces

A. Representation in metagenomes from different environments
B. Association with different potential hosts

**Supplementary Figure 3**

Multiple alignment of the sequences of homologs of disulfide-bonded, myristoylated proteins G9/A16

**Supplementary Figure 4**

Multiple alignment of the sequences of homologs of disulfide-bonded, myristoylated proteins L1/F9.

**Supplementary Figure 5**

Structural model of the SmDNAV ssDNA-binding protein

**Supplementary Figure 6**

Relative phylogenetic depths of nucleocytoviricot orders and families

## Notes

### Competing Interest Statement

The authors have declared no competing interest.

