## Supplementary figures for "Loss of replication and transcription systems accompanying transition to nucleus-dependent replication in “Ariadnavirales”, a proposed new order in nucleocytoviricot class *Megaviricetes*"

Fig. S1

MCP

(cls006)

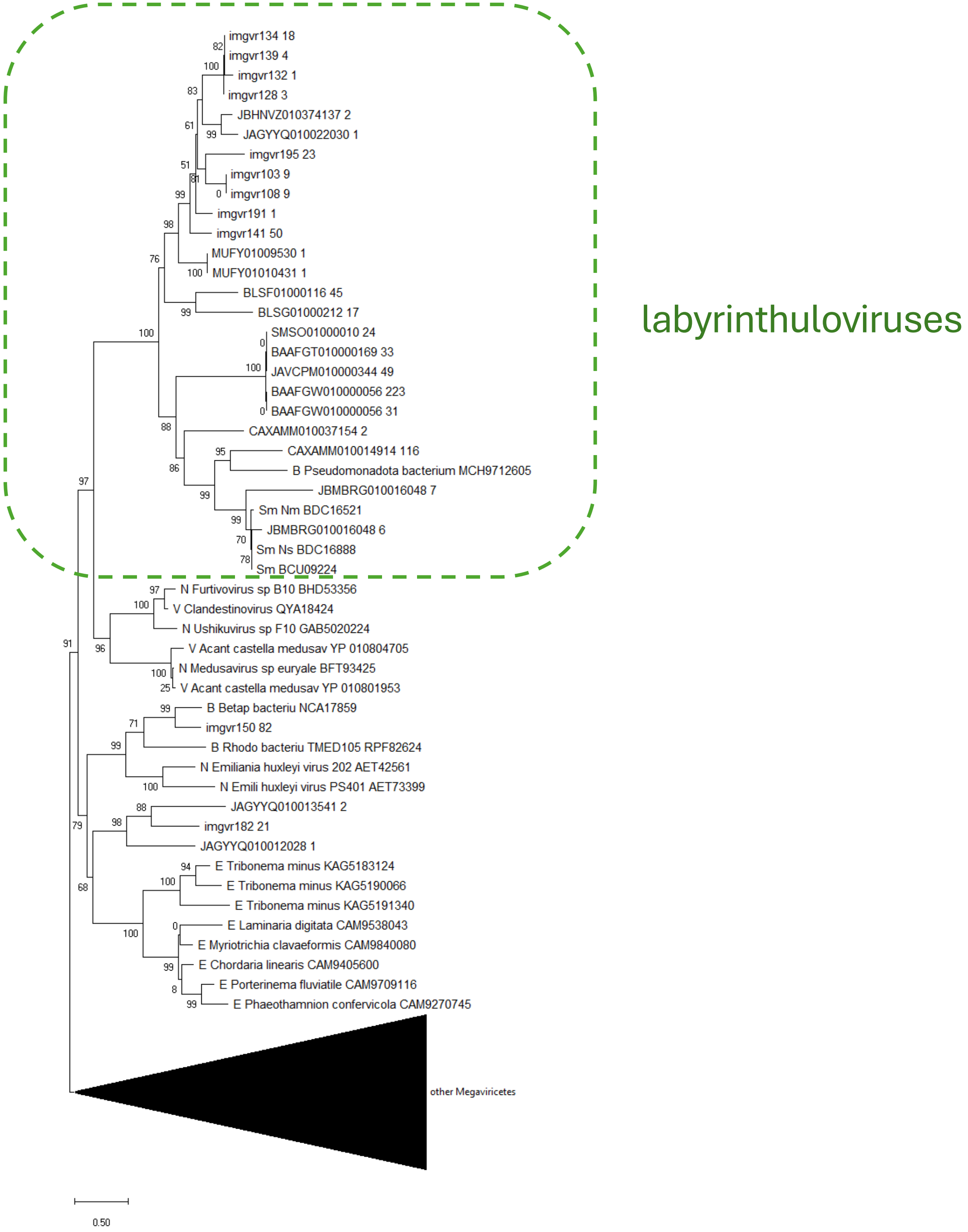

Fig. S2

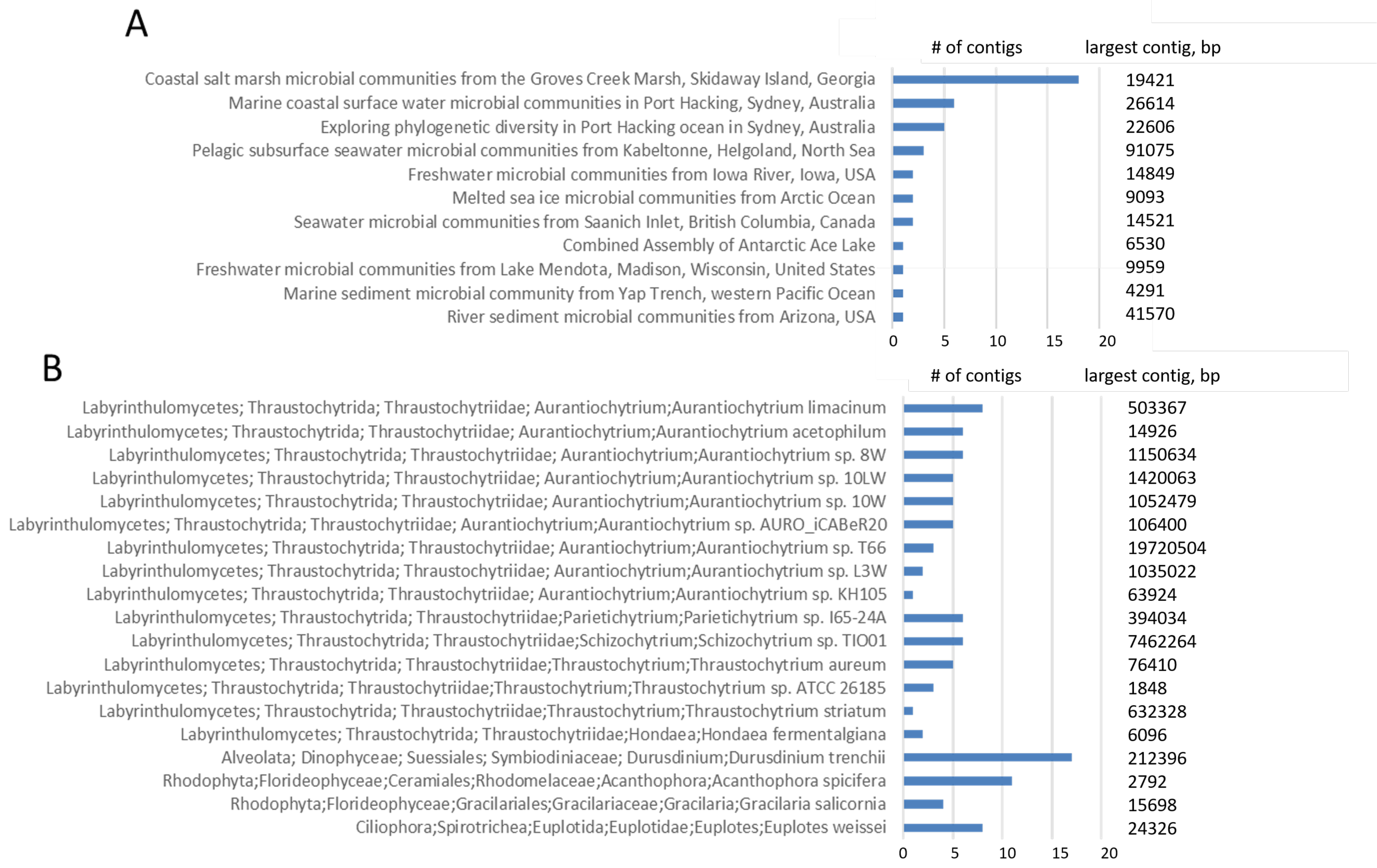
A
### of contigs
largest contig, bp
Coastal salt marsh microbial communities from the Groves Creek Marsh, Skidaway Island, Georgia
Marine coastal surface water microbial communities in Port Hacking, Sydney, Australia
Exploring phylogenetic diversity in Port Hacking ocean in Sydney, Australia
Pelagic subsurface seawater microbial communities from Kabeltonne, Helgoland, North Sea
Freshwater microbial communities from Iowa River, Iowa, USA
Melted sea ice microbial communities from Arctic Ocean
Seawater microbial communities from Saanich Inlet, British Columbia, Canada
Combined Assembly of Antarctic Ace Lake
Freshwater microbial communities from Lake Mendota, Madison, Wisconsin, United States
Marine sediment microbial community from Yap Trench, western Pacific Ocean
River sediment microbial communities from Arizona, USA
19421
26614
22606
91075
14849
9093
14521
6530
9959
4291
41570
0
5
10
15
20
B
### of contigs
largest contig, bp
Labyrinthulomycetes; Thraustochytrida; Thraustochytriidae; Aurantiochytrium;Aurantiochytrium limacinum
Labyrinthulomycetes; Thraustochytrida; Thraustochytriidae; Aurantiochytrium;Aurantiochytrium acetophilum
Labyrinthulomycetes; Thraustochytrida; Thraustochytriidae; Aurantiochytrium;Aurantiochytrium sp. 8W
Labyrinthulomycetes; Thraustochytrida; Thraustochytriidae; Aurantiochytrium;Aurantiochytrium sp. 10LW
Labyrinthulomycetes; Thraustochytrida; Thraustochytriidae; Aurantiochytrium;Aurantiochytrium sp. 10W
Labyrinthulomycetes; Thraustochytrida; Thraustochytriidae; Aurantiochytrium;Aurantiochytrium sp. AURO_iCABeR20
Labyrinthulomycetes; Thraustochytrida; Thraustochytriidae; Aurantiochytrium;Aurantiochytrium sp. T66
Labyrinthulomycetes; Thraustochytrida; Thraustochytriidae; Aurantiochytrium;Aurantiochytrium sp. L3W
Labyrinthulomycetes; Thraustochytrida; Thraustochytriidae; Aurantiochytrium;Aurantiochytrium sp. KH105
Labyrinthulomycetes; Thraustochytrida; Thraustochytriidae;Parietichytrium;Parietichytrium sp. I65-24A
Labyrinthulomycetes; Thraustochytrida; Thraustochytriidae;Schizochytrium;Schizochytrium sp. TIO01
Labyrinthulomycetes; Thraustochytrida; Thraustochytriidae;Thraustochytrium;Thraustochytrium aureum
Labyrinthulomycetes; Thraustochytrida; Thraustochytriidae;Thraustochytrium;Thraustochytrium sp. ATCC 26185
Labyrinthulomycetes; Thraustochytrida; Thraustochytriidae;Thraustochytrium;Thraustochytrium striatum
Labyrinthulomycetes; Thraustochytrida; Thraustochytriidae;Hondaea;Hondaea fermentalgiana
Alveolata; Dinophyceae; Suessiales; Symbiodiniaceae; Durusdinium;Durusdinium trenchii
Rhodophyta;Florideophyceae;Ceramiales;Rhodomelaceae;Acanthophora;Acanthophora spicifera
Rhodophyta;Florideophyceae;Gracilariales;Gracilariaceae;Gracilaria;Gracilaria salicornia
Ciliophora;Spirotrichea;Euplotida;Euplotidae;Euplotes;Euplotes weissei
503367
14926
1150634
1420063
1052479
106400
19720504
1035022
63924
394034
7462264
76410
1848
632328
6096
212396
2792
15698
24326
0
5
10
15
20

### Fig. S3

#### cls_004 Myristoylated protein G9/A16 homolog

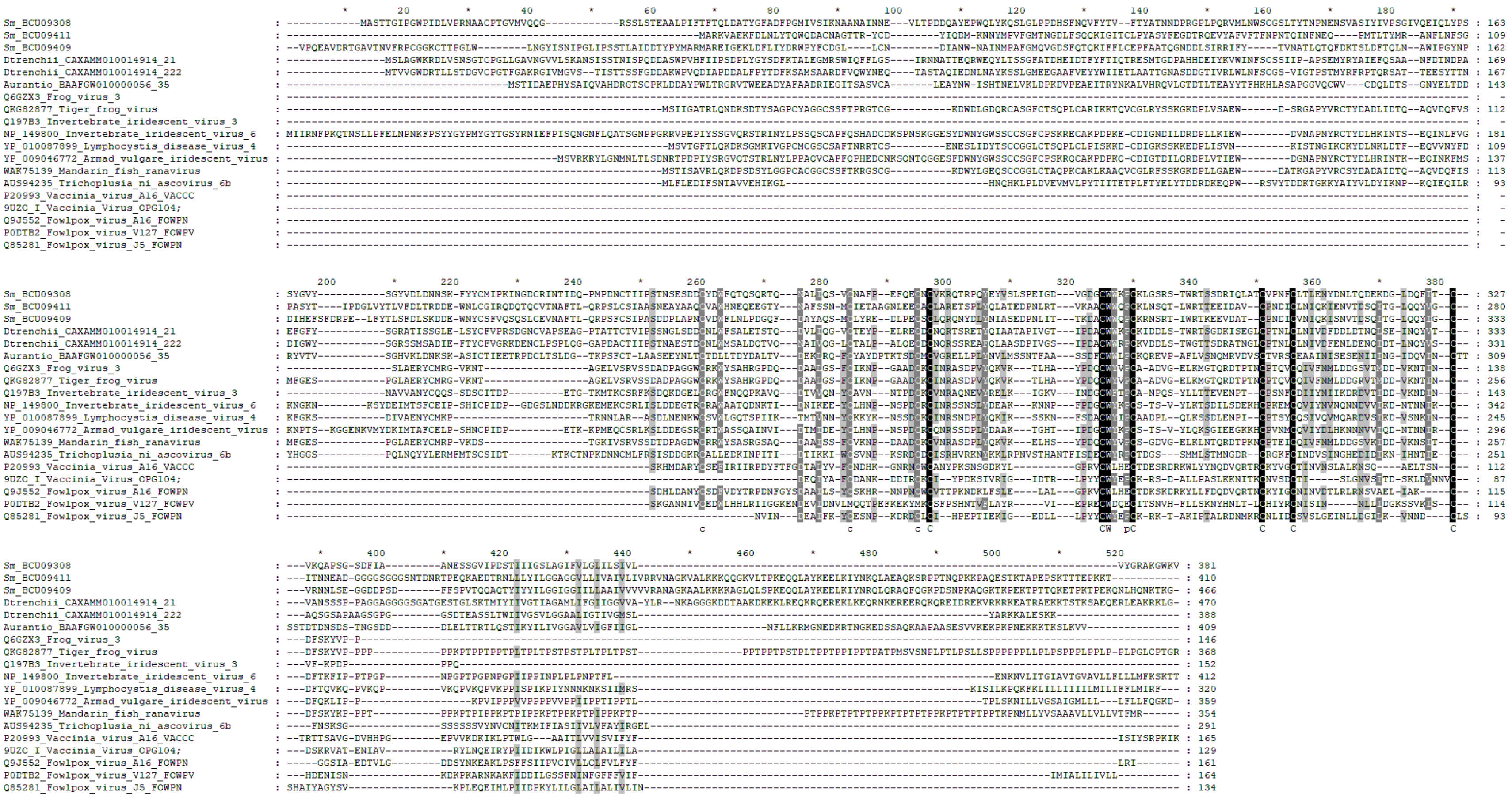

cls_004_for_figureS3.afa

### Fig. S4

#### cls_034 Myristoylated protein L1R/F9L homolog

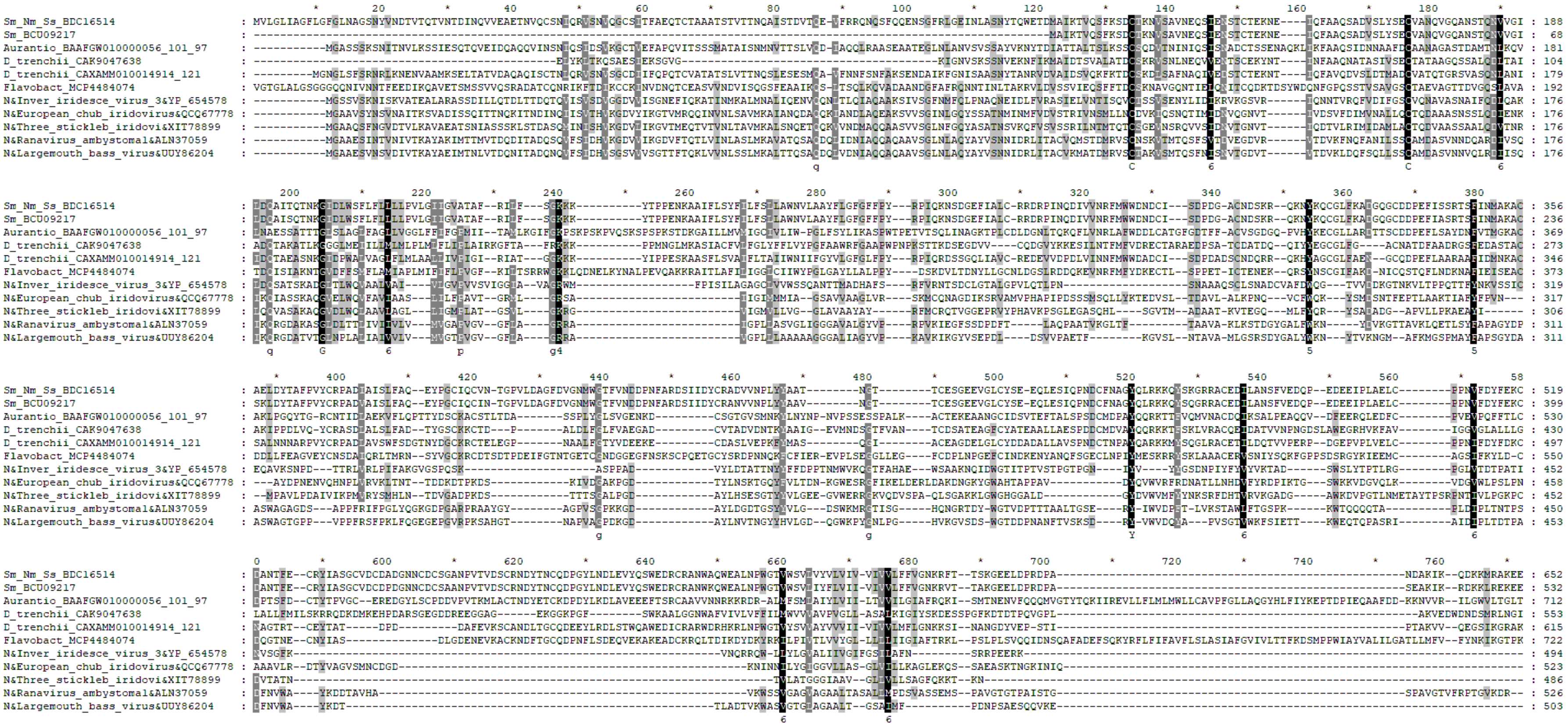

cls_034_for_figure.afa

Fig. S5

ssDNA-binding protein

cls028

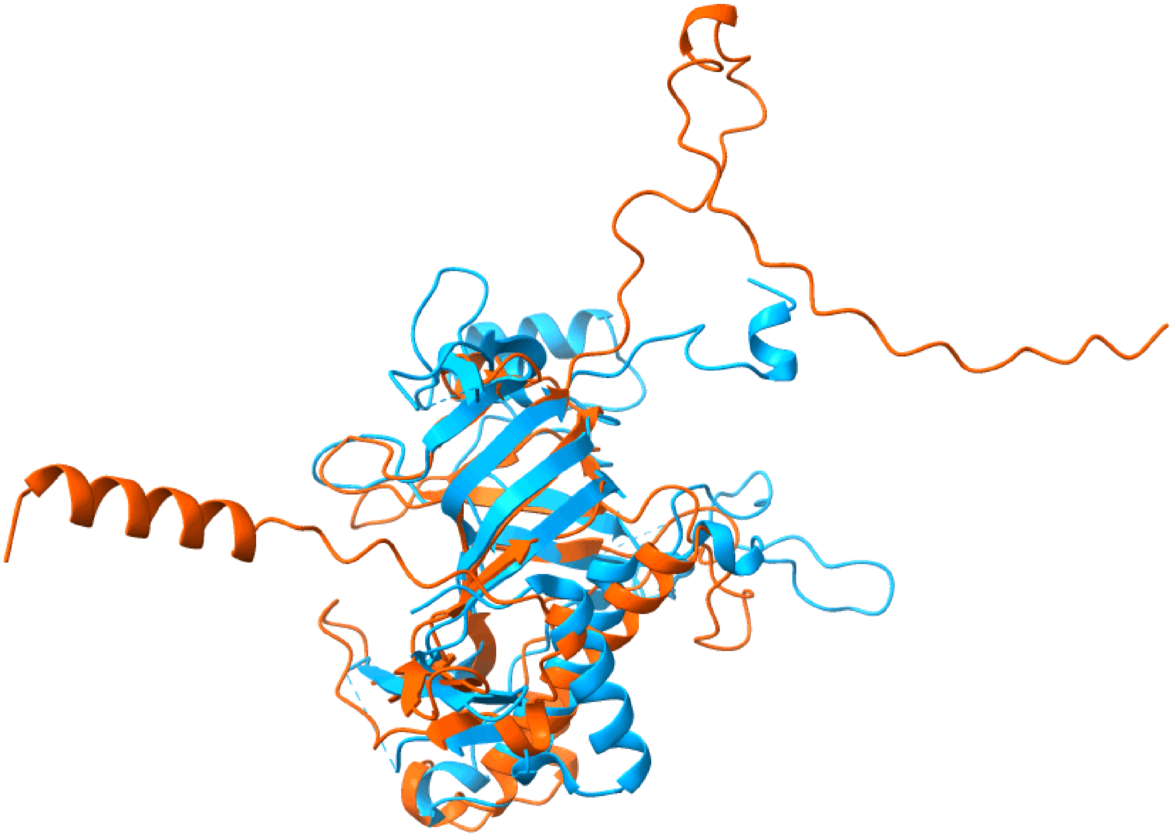

7yeq_A
cls_028/JAAEPP010000199_18/BCU09318.1
plddt: 0.8888
FoldSeek evalue: $6.547 \times 10^{-8}$

Fig. S6

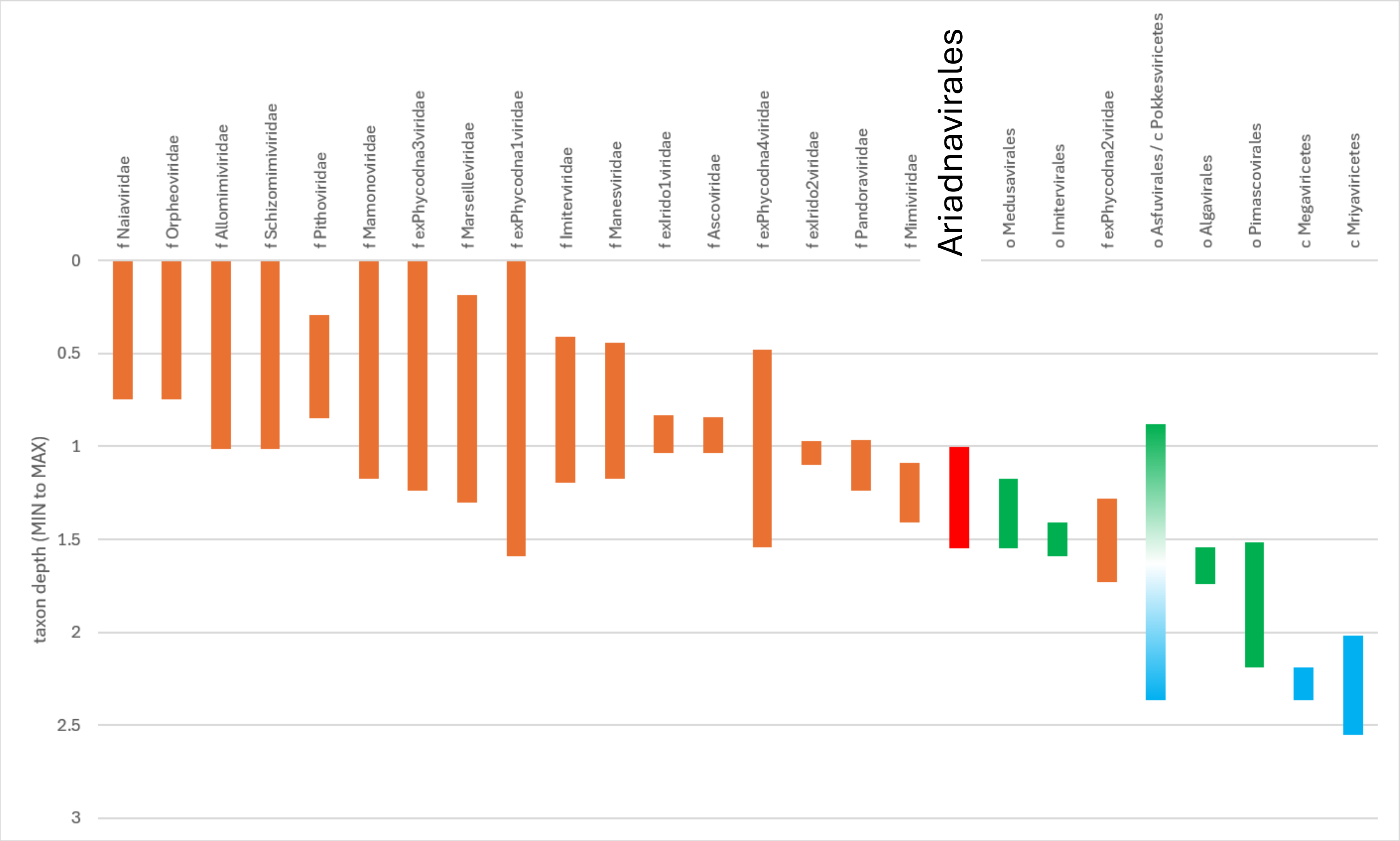

taxon depth (MIN to MAX)
0
0.5
1
1.5
2
2.5
3
f Naiaviridae
f Orpheoviridae
f Allomimiviridae
f Schizomimiviridae
f Pithoviridae
f Mamonoviridae
f exPhycodna3viridae
f Marseilleviridae
f exPhycodna1viridae
f Imitervirdae
f Manesviridae
f exIrido1viridae
f Ascoviridae
f exPhycodna4viridae
f exIrido2viridae
f Pandoraviridae
f Mimiviridae
Ariadnavirales
o Medusavirales
o Imitervirales
f exPhycodna2viridae
o Asfuvirales / c Pokkesviricetes
o Algavirales
o Pimascovirales
c Megaviricetes
c Mriyaviricetes
